# Analysis of culture and RNA isolation methods for precision-cut liver slices from cirrhotic rats

**DOI:** 10.1101/2023.07.18.549535

**Authors:** Ben D. Leaker, Yongtao Wang, Joshua Tam, R. Rox Anderson

## Abstract

Precision-cut liver slices (PCLS) are increasingly used as a model to investigate anti-fibrotic therapies. However, many studies use PCLS from healthy animals treated with pro-fibrotic stimuli in culture, which reflects only the early stages of fibrosis. The performance of PCLS from cirrhotic animals has not been well characterized and there is no consensus on optimal culture conditions. In this study, we report a method for the collection and culture of cirrhotic PCLS and compare the effect of common culture conditions on viability, function, and gene expression. Additionally, we compared three methods of RNA isolation and identified a protocol with high yield and purity. We observed significantly increased albumin production when cultured with insulin and dexamethasone, and when incubated on a rocking platform. Culturing with insulin and dexamethasone maintained gene expression closer to the levels in fresh slices. However, despite stable viability and function up to 4 days, we found significant changes in expression by day 2. Due to the influence of matrix stiffness on fibrosis and hepatocellular function, it is important to evaluate prospective anti-fibrotic therapies in a platform that preserves tissue biomechanics. PCLS from cirrhotic animals represent a promising tool for the development of treatments for chronic liver disease.

## Introduction

Conventional cell culture systems lack the complexity of tissues, while preclinical animal models typically have limited throughput. In contrast, organ or tissue culture methods preserve complexity while allowing assays similar to those performed in cell cultures. Precision-cut liver slices (PCLS) is an ex vivo tissue culture model that is gaining popularity for the study of liver fibrosis. PCLS preserve the extracellular matrix and resident liver cell populations, providing a more accurate representation of the in vivo interactions between different cell types, and between cells and the matrix. PCLS also enable higher throughput and more easily controlled experiments than in vivo models as dozens of slices can be obtained from one animal. The primary drawbacks of PCLS are the limited time they can be cultured – typically a few days – as well as changes in expression that occur during culture.

PCLS have particularly attracted interest as a platform to test anti-fibrotic therapies. Many previous experiments have used healthy PCLS treated with pro-fibrotic stimuli, such as TGFβ and PDGF, to induce fibrosis ex vivo^1-9^, or have relied on the spontaneous fibrosis activated in PCLS in culture^10-16^. Such experiments are useful to study the earliest onset of liver fibrosis but do not capture many of the effects of chronic liver disease in clinically relevant liver fibrosis. Chronic liver injury induces senescence of a variety of liver cell populations^17,18^, and the long-term deposition of collagen and associated increase in tissue stiffness can suppress hepatocellular function through HNF4a and cytochrome P450^19-21^. There is also a critical positive feedback loop between fibrosis and matrix stiffness^22-25^, which could significantly influence the effects of prospective anti-fibrotic therapies.

For a more clinically relevant model, PCLS can be collected from patient biopsies or from animal models after liver fibrosis or cirrhosis is induced in vivo. A variety of fibrosis etiologies have been investigated with this approach, including alcohol^26^, hepatitis infection^27^, non-alcoholic steatohepatitis (NASH)^26,28^, cholestasis^29-33^, and chemical hepatotoxins^26,34,35^.

The performance of PCLS from cirrhotic animals in culture has not been well characterized. Consequently, prior work with fibrotic and cirrhotic slices often reference the methods of experiments performed with slices from healthy animals, which may not be applicable given the significant differences between the healthy and cirrhotic liver. There is also no consensus on appropriate culture conditions, most notably whether to include insulin and dexamethasone in the culture media. A direct comparison of the effect of culture conditions on viability, function, and gene expression has not previously been performed.

To move towards standardized methodology, we report a method for the collection and culture of PCLS from cirrhotic rats. We compare the effect of culture media and rocking on PCLS viability and albumin production, as well as characterize changes in expression of key genes over time and when cultured with and without insulin and dexamethasone. We also compare three methods for the isolation of RNA from cirrhotic PCLS and report a method capable of obtaining RNA with higher yield and purity than the standard RNeasy and TRIzol protocols.

## Methods

### Animals

All animal work was approved by the Massachusetts General Hospital Institutional Animal Care and Use Committee. All experiments were performed in accordance with relevant guidelines and regulations. Male Wistar rats were purchased from Charles River Laboratories. Animals were housed in a controlled environment with food and water ad libitum. Cirrhosis was induced with biweekly intraperitoneal injections of thioacetamide at 200mg/kg for 12 weeks followed by a 1 week wash out period.

### PCLS collection

Animals were euthanized via cardiac puncture under sterile conditions. The whole liver was harvested and 8mm biopsy punches were used to obtain tissue columns. Columns were collected from the thickest portions of the medial and left lobe. The columns were immediately placed in chilled sterile Krebs Henseleit buffer (KHB; Sigma Aldrich, USA).

Columns were cut in half with a scalpel to obtain 2 shorter columns (Fig. 1a). These columns were mounted to the platform of a 7000smz-2 vibratome (Campden Instruments Limited, UK) with cyanoacrylate glue and the cutting tray of the vibratome was filled with chilled sterile KHB (Fig. 1b). The vibratome was fitted with a ceramic blade (Campden Instruments Limited) and run at 50Hz frequency, 2.5mm amplitude, 0.15cm/s advance speed, 250μm slice thickness (Fig. 1c). The first few slices were discarded to ensure a flat tissue surface. After cutting, slices were transferred to a sterile dish with chilled sterile KHB. A 6mm biopsy punch was used to trim the final PCLS. This step ensures all PCLS are uniform in size, as the initial columns are often irregularly shaped (Fig. 1d). With such small samples, these irregular edges can result in significant differences in the total amount of tissue. This step also removes any tissue that may have come into contact with glue. The trimmed PCLS were then transferred to a 24 well plate with chilled sterile KHB. The total time from harvesting of the liver to the end of collecting PCLS was around 1 hour. Approximately 20 uniform, viable slices were obtained per animal.

**Figure 1:**
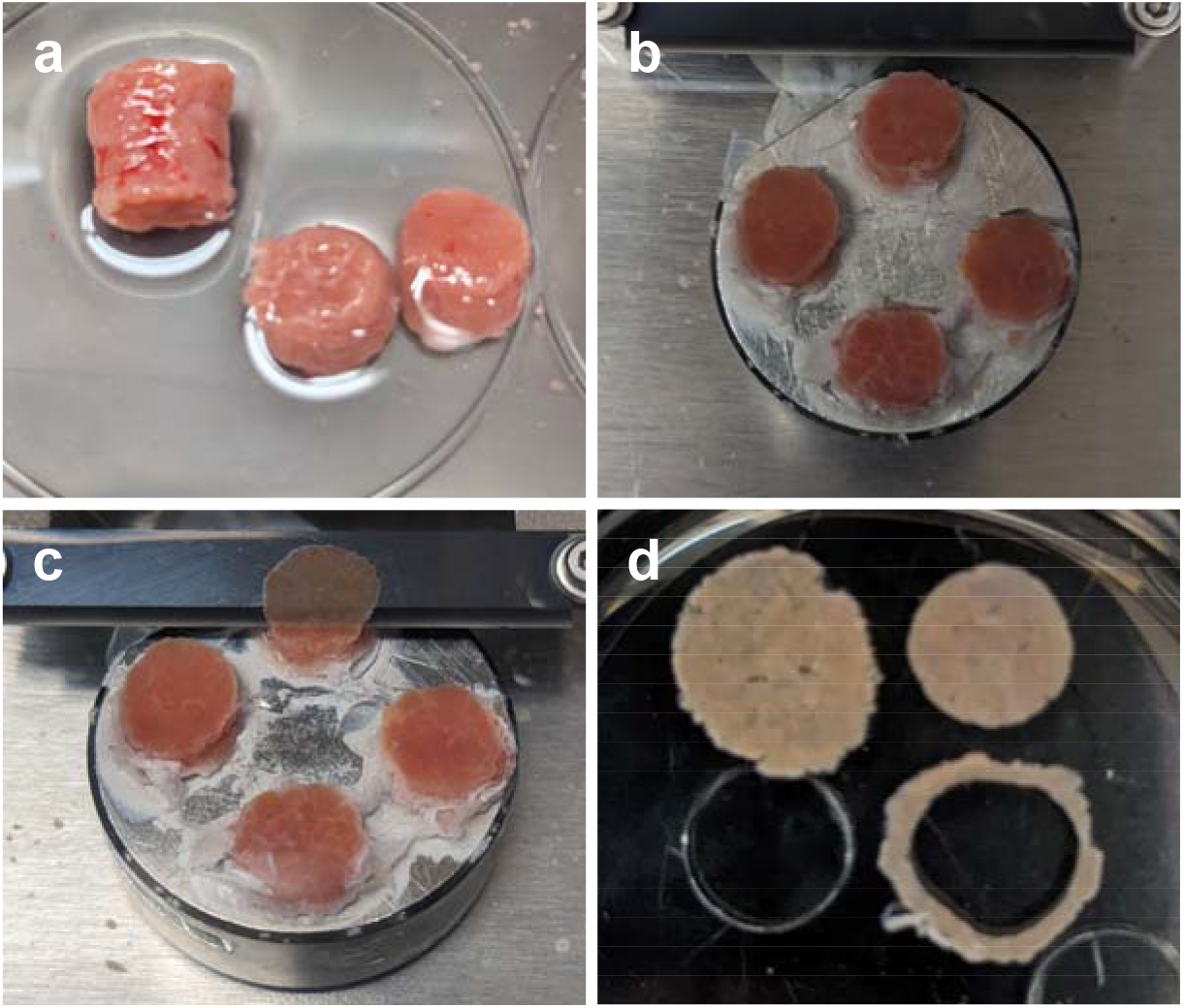
PCLS collection. (A) Tissue columns were harvested with an 8mm biopsy punch then cut into 2 shorter columns. (B) Columns were mounted to the vibratome platform with cyanoacrylate glue. (C) 250μm thick PCLS were cut at 50Hz, 2.5mm amplitude, 0.15cm/s. (D) PCLS were trimmed with a 6mm biopsy punch to ensure uniform slices.

### PCLS culture

PCLS were placed on an 8μm-pore Transwell insert (Corning, USA) in a 12 well plate. All slices were cultured in 1mL Williams’ Medium E (Sigma Aldrich) with 2% FBS, 1% pen-strep, and 1% L-glutamine (ThermoFisher, USA). Some slices were additionally cultured with 100nM dexamethasone (Sigma Aldrich) and 1X insulin-transferrin-selenium-ethanolamine (ThermoFisher). Plates were incubated under standard cell culture conditions either stationary or on a rocking platform. Media was changed every 24hrs.

### MTS viability assay

PCLS were placed in a 48 well plate with 400μL culture media and 80μL MTS reagent (Abcam, USA). The plates were incubated on a rocking platform under standard cell culture conditions for 1hr then the media was collected and absorbance was measured at 490nm. Measurements were normalized to the viability of fresh slices.

### Albumin ELISA

Media collected from the culture plate was centrifuged at 4°C, 400g for 10min. The supernatant was then stored at -20°C until ready to use. Albumin concentration was measured with a rat albumin ELISA kit (Abcam) according to manufacturer instructions.

### RNA Isolation & qPCR

3 methods of RNA isolation were compared. The first method used the RNeasy micro kit (Qiagen, Germany) according to manufacturer instructions. The second method used TRIzol (Invitrogen, USA) according to the manufacturer instructions. The final method was a protocol reported by Ziros et al. ^36^ for the isolation of RNA from mouse thyroid, a very small organ which requires high yield RNA extraction. Briefly, PCLS were homogenized in TRIzol and frozen at -80°C for at least 30min. Phase separation was performed with 1-bromo-3-chloropropane (Sigma Aldrich) and the aqueous phase was purified using RNeasy micro spin columns. For all methods, PCLS were homogenized with TissueLyser II (Qiagen).

RNA concentration and purity were measured with NanoDrop (ThermoFisher). RNA integrity was confirmed by gel electrophoresis on E-gel EX 2% agarose gels (Thermofisher).

cDNA was prepared from 1μg total RNA using the SuperScript VILO IV kit (Thermofisher) according to manufacturer instructions. qPCR was then performed with TaqMan Fast Advanced Master Mix (Thermofisher) and TaqMan fluorescent probes for HNF4A (Thermofisher; assay ID: Rn04339144_m1), CYP1A2 (Rn00561082_m1), LYVE1 (Rn01510421_m1), TGFB1 (Rn00572010_m1), COL1A1 (Rn01463848_m1), ACTA2 (Rn01759928_m1), CDKN1A (Rn00589996_m1), and MKI67 (Rn01451446_m1).

Stability analysis was performed with NormFinder^37^ to determine appropriate reference genes. Fresh slices and PCLS after 4 days in culture were compared with TaqMan probes for B2M (Rn00560865_m1), YWHAZ (Rn00755072_m1), HPRT1 (Rn01527840_m1), HMBS (Rn01421873_g1), 18S (Hs99999901_s1), SDHA (Rn00590475_m1), and UBC (Rn01789812_g1). HPRT1 and HMBS were found to be the best combination of reference genes, with a stability value of 0.063.

### Statistics

Student’s t-test was performed to assess statistical significance with a p-value threshold of 0.05. The Benjamini-Hochberg procedure was used to correct for multiple comparisons in the qPCR experiments. Yield and purity metrics are expressed as mean ± standard deviation.

## Results

### Effect of culture conditions on cirrhotic PCLS viability and function

Two parameters were evaluated for their effect on cirrhotic PCLS viability and function: media supplements (insulin-transferrin-selenium-ethanolamine and dexamethasone) and incubation on a rocking platform. PCLS cultured with these supplements had no significant difference in overall viability after 4 days in culture (Fig. 2A) but the supplemented media significantly increased PCLS function at all timepoints (Fig. 2B). Similarly, incubation on a rocking platform did not have an effect on the overall viability (Fig. 2C) but preserved PCLS function at later timepoints (Fig. 2D).

**Figure 2:**
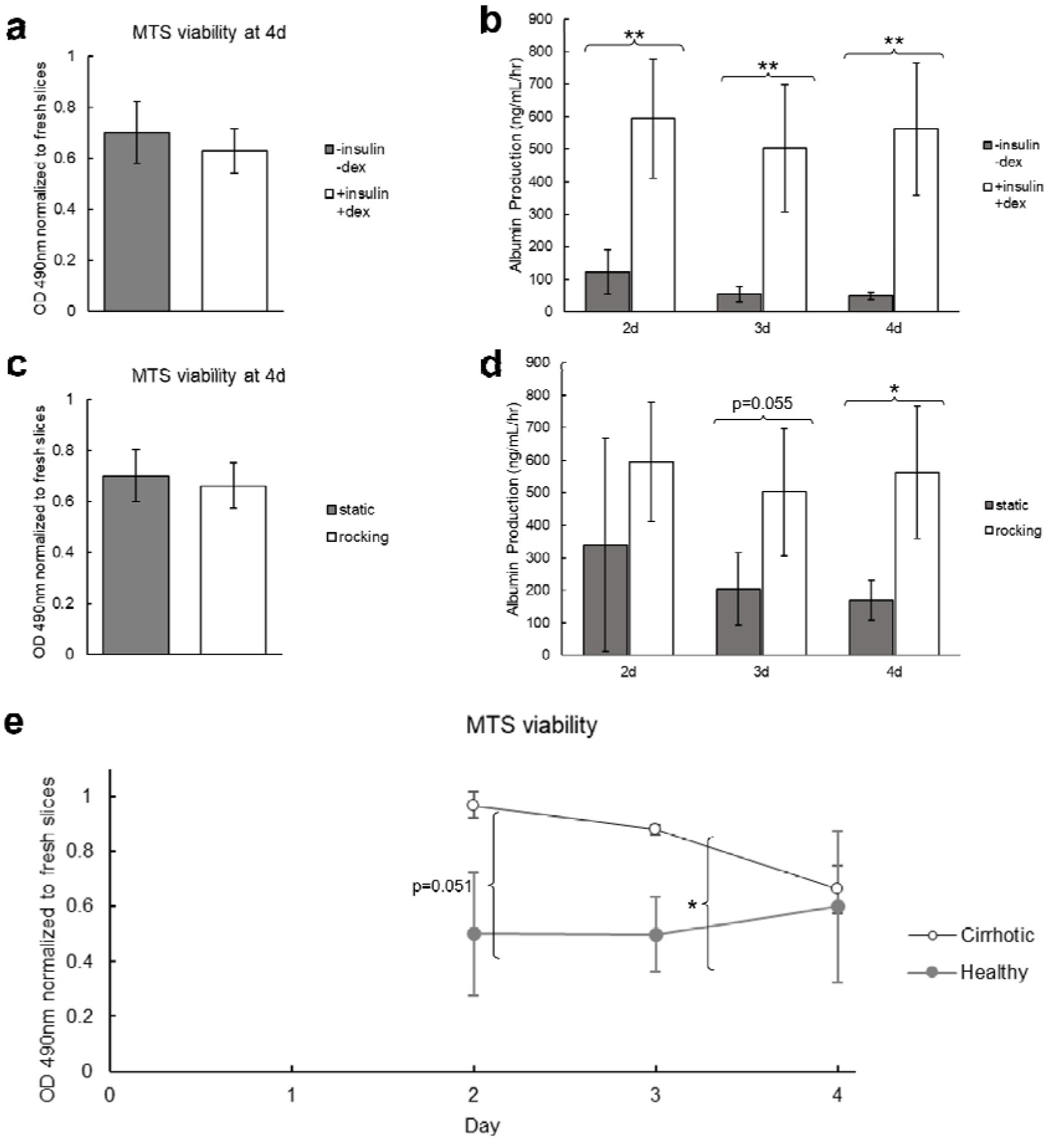
Comparison of culture conditions for cirrhotic PCLS viability and function. (A&B) Cirrhotic PCLS viability and function when cultured with and without insulin-transferrin-selenium-ethanolamine and dexamethasone. Both groups were incubated on a rocking platform. (A) Viability measured by MTS assay after 4 days in culture normalized to the viability of fresh PCLS. There is no significant difference in slice viability. (B) ELISA for albumin secreted into the media over 4 days in culture. Culturing with insulin-transferrin-selenium-ethanolamine and dexamethasone significantly increases PCLS function at all timepoints. (C&D) Cirrhotic PCLS viability and function when incubated on a rocking platform or static. Both groups were cultured with insulin-transferrin-selenium-ethanolamine and dexamethasone. (C) Viability measured by MTS assay after 4 days in culture. There is no significant difference in slice viability (D) ELISA for albumin secreted into the media over 4 days in culture. Culturing on a rocking platform significantly increased PCLS function at the 4 day timepoint. (E) Viability of healthy and cirrhotic PCLS measured by MTS assay over 4 days in culture normalized to viability of fresh healthy and cirrhotic PCLS, respectively. The cirrhotic slices maintain higher viability until day 4. * p<0.05, **p<0.01

Interestingly, we also found that cirrhotic PCLS maintained higher viability in culture than PCLS from healthy animals until day 4 (Fig. 2E). This may be related to the poor perfusion and hypoxic environment of the cirrhotic liver in vivo ^38-41^. Survival in culture is heavily dependent on oxygen delivery to the tissue, so it is possible that adaptation to hypoxia in vivo enables better survival ex vivo.

### Comparison of RNA isolation methods for cirrhotic PCLS

RNA yield and purity for three isolation methods – Qiagen RNeasy kit, standard TRIzol protocol, and the small tissue RNA isolation protocol described by Ziros et al. for the isolation of high-quality RNA from mouse thyroid – were compared with cirrhotic PCLS after 4 days in culture (Table 1). The RNeasy protocol produced high purity RNA, but with very low yield. Conversely, the TRIzol protocol gave higher yield but with poor purity. The protocol described by Ziros et al. gave the highest yield and purity of all 3 protocols. To determine whether the difference in purity would significantly alter expression measurements, RNA isolated by the TRIzol and Ziros et al. protocols were compared with qPCR. Of the 8 genes measured, 1 was found to have a statistically significant difference in expression after multiple hypothesis correction (Supplementary Figure 1). We also found that there is a significant decrease in RNA yield from PCLS in culture compared to fresh slices in addition to a small but statistically significant decrease in A^260^/^280^ (Supplementary Table 1).

**Table 1:** Yield and purity metrics for three methods of RNA isolation from cirrhotic PCLS after 4 days in culture. The protocol described by Ziros et al. gives the highest yield and RNA purity. *p<0.05 comparing Ziros et al. and Qiagen RNeasy protocols, ^†^p<0.05 comparing Ziros et al. and Trizol

| RNA Isolation Protocol | Yield (μg/mg tissue) | A | A |
| --- | --- | --- | --- |
| Qiagen RNeasy | 0.2±0.1 | 2.01±0.03 | 1.7±0.4 |
| TRIzol | 1.5±0.1 | 1.89±0.03 | 0.6±0.1 |
| Ziros et al. | 2.3±0.9* | 2.03±0.03† | 1.9±0.4† |

### Effect of culture media on expression in cirrhotic PCLS

Expression of 8 key genes related to liver function and fibrosis were evaluated by qPCR for cirrhotic slices cultured for 4 days with and without insulin-transferrin-selenium-ethanolamine and dexamethasone (Fig. 3). In all cases with statistically significant differences, culturing with these supplements resulted in expression more similar to fresh slices. HNF4A and CYP1A2, markers of hepatocyte function and differentiation, were increased with the supplemented media. TGFB1, a key gene associated with the spontaneous ex vivo fibrosis response, was reduced with the supplemented media, as well as MKI67, a marker of proliferation. There were no statistically significant differences in expression of LYVE1, a marker of liver sinusoidal endothelium, COL1A1 and ACTA2, markers of fibrosis, or CDKN1A (p21), a marker of senescence.

**Figure 3:**
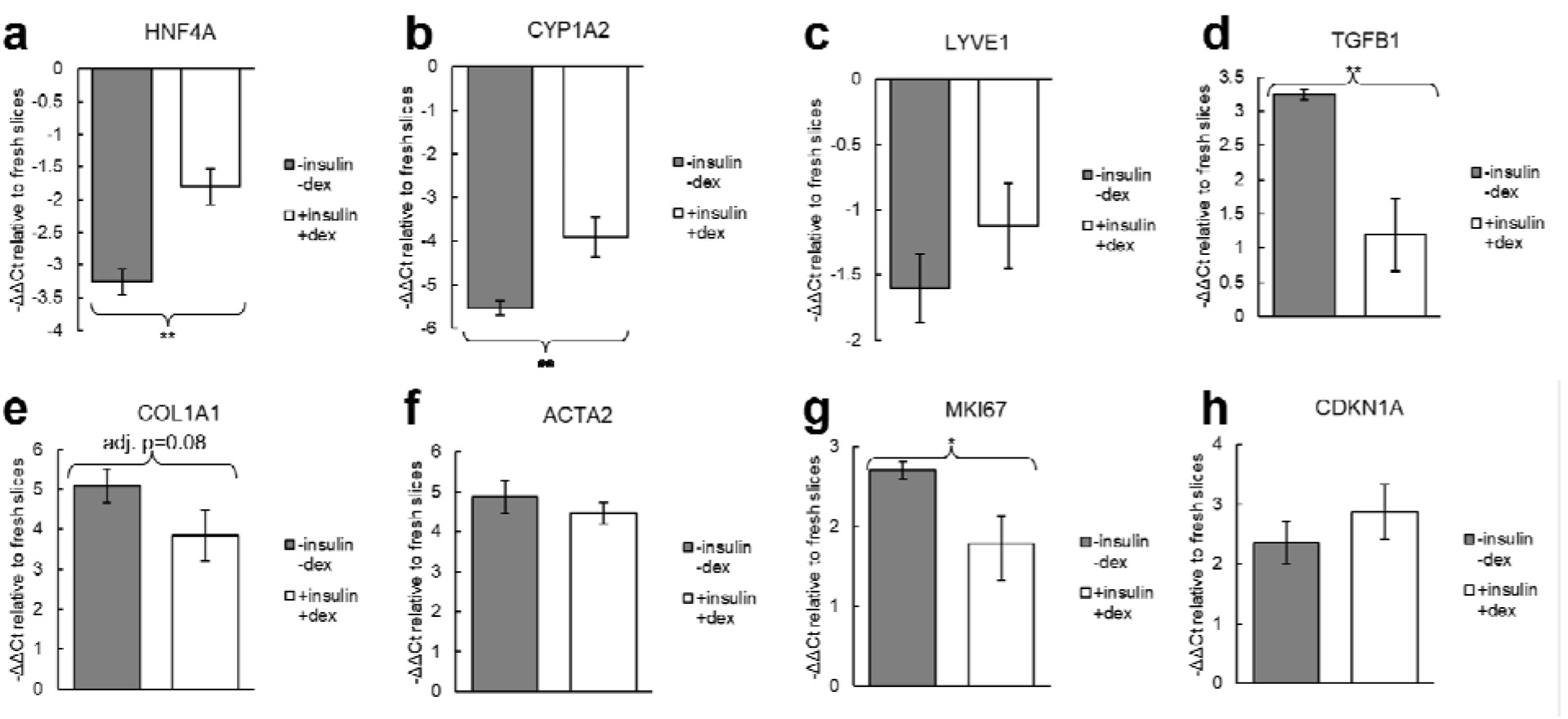
Expression changes in cirrhotic PCLS after 4 days in culture with and without insulin-transferrin-selenium-ethanolamine and dexamethasone. Both groups were incubated on a rocking platform. ΔΔCt values were calculated relative to fresh slices. For all genes with statistically significant differences, culturing with insulin-transferrin-selenium-ethanolamine and dexamethasone maintained expression closer to fresh slices. Statistical tests were performed on the ΔCt values. *p<0.05, **p<0.01

### Time course of expression changes in cirrhotic PCLS

Expression of these 8 genes were measured at the 2 day and 4 day timepoints to characterize how expression changes over time in culture (Fig. 4). For these results, all slices were cultured with insulin-transferrin-selenium-ethanolamine and dexamethasone, and were incubated on a rocking platform. We found that despite stable viability and function, there are already significant differences in many genes by the 2 day timepoint. HNF4A and CYP1A2 are significantly downregulated compared to fresh slices by day 2, and expression further decreases by day 4. LYVE1 expression is only significantly decreased at the 4 day timepoint. Significantly elevated expression of TGFB1, COL1A1, and ACTA2 is not observed until the 4 day timepoint, indicating that the ex vivo spontaneous pro-fibrotic stimuli induce a relatively slow process in cirrhotic PCLS. CDKN1A expression is significantly elevated by day 2 and there is no significant difference in expression between the 2 day and 4 day timepoints. Interestingly, MKI67 expression is decreased at the 2 day timepoint but increased by day 4.

**Figure 4:**
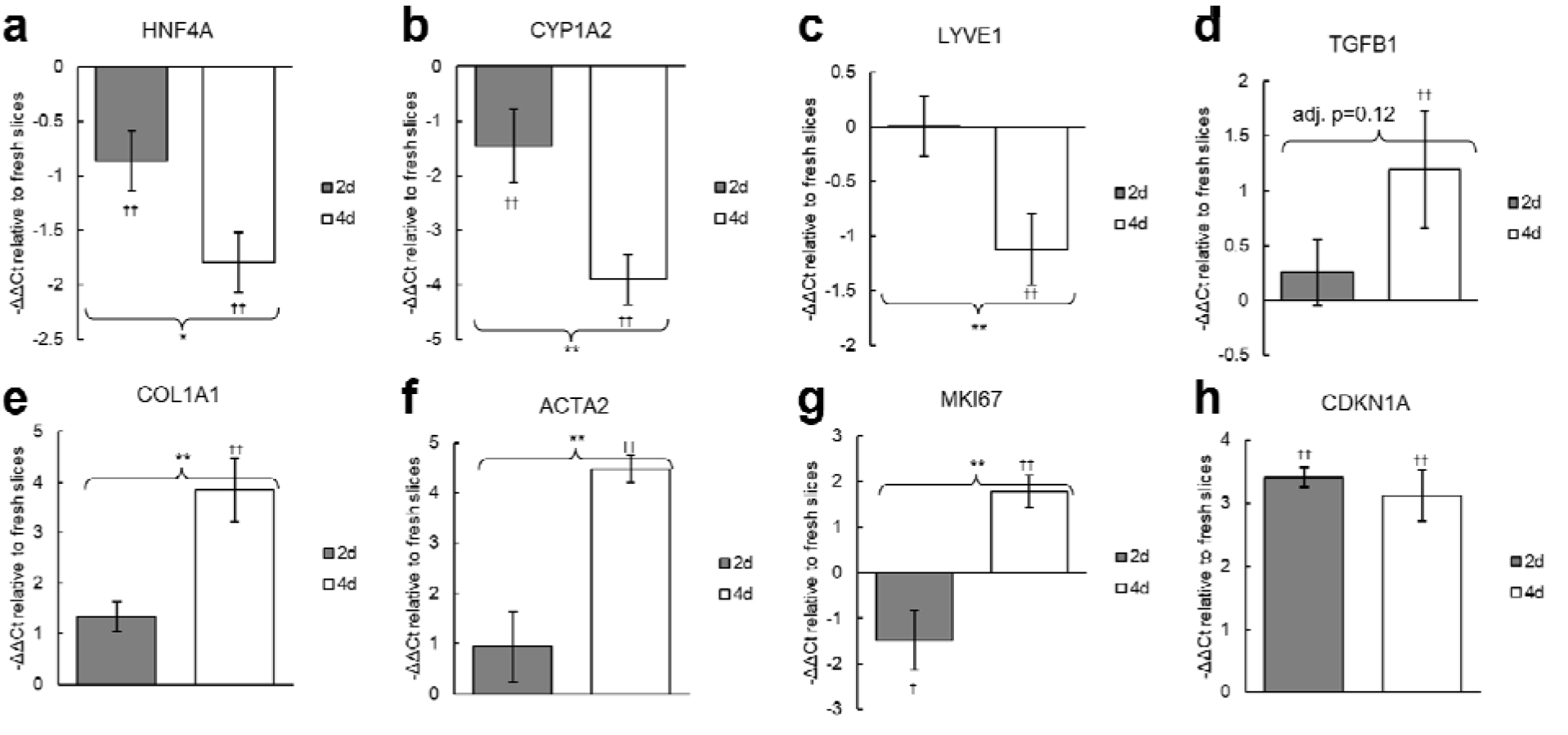
Expression changes in cirrhotic PCLS at 2 days and 4 days in culture. Slices were cultured with insulin-transferrin-selenium-ethanolamine and dexamethasone, and were incubated on a rocking platform. ΔΔCt values were calculated relative to fresh slices. Statistical tests were performed on the ΔCt values. *p<0.05, **p<0.01; ^†^p<0.05 compared to fresh slices, ^††^p<0.01 compared to fresh slices.

## Discussion

In this work we characterized the performance of cirrhotic PCLS in culture and clarified the effect of different commonly used culture conditions. We describe a method for the collection and culture of cirrhotic PCLS that are viable with stable function for up to 4 days. Our results show that there is a clear benefit to culturing with insulin and dexamethasone as well as incubating on a rocking platform. This method significantly improved albumin production and maintained expression of HNF4A, CYP1A2, TGFB1, and MKI67 closer to the level in fresh slices. We would recommend using these methods unless they are contraindicated by the specific hypothesis under investigation (e.g., insulin interfering with a signaling pathway of interest).

Viability and albumin production are the most commonly used methods in prior work to characterize the performance of PCLS in culture. However, we also investigated changes in expression of key liver genes and found several important changes that are not reflected by these assays. Despite high albumin production throughout the time in culture, there was progressive decline in expression of HNF4A and CYP1A2, two key regulators of hepatocellular function. This is possibly due to the same process of dedifferentiation that is commonly observed with primary hepatocytes in culture^42-46^. We also found that the senescence marker CDKN1A is upregulated at both day 2 and 4. Hepatocyte senescence is associated with a pro-inflammatory secretory state, declining function, and suppressed proliferation^18,47^. Interestingly, we observed that the proliferation marker MKI67 is downregulated at the 2 day timepoint, but becomes upregulated by day 4. This biphasic response may indicate differing proliferative states for different cell populations. It is possible that hepatocyte senescence drives MKI67 expression down at day 2, while a delayed fibrotic response – demonstrated by upregulation of TGFB1, COL1A1, and ACTA2 – activates proliferation of stellate cells at the 4 day timepoint. We also found that LYVE1 is downregulated at day 4. LYVE1 is a marker of liver sinusoidal endothelial cells, and its downregulation is associated with capillarization of sinusoids^48,49^. This may indicate some damage or dedifferentiation is occurring to the sinusoidal endothelium in culture.

These results demonstrate that different metrics provide different results for the stability of PCLS in culture. Viability and albumin production remain high up to 4 days in culture. However, when compared to fresh slices there are significant changes in expression of key genes as early as day 2. It is difficult to reconcile these data to produce a single number for the time cirrhotic PCLS can be cultured. An appropriate timeline needs to be determined on a case-by-case basis depending on the specific hypothesis and metrics under investigation.

We have also reported a protocol for RNA isolation with higher yield and purity than the standard RNeasy and TRIzol protocols. High yield is necessary to extract sufficient RNA for quality control and downstream assays from small tissue samples, while high purity is required to prevent contaminants from interfering in expression measurements. The general guideline for pure RNA is A^260^/_230_ around 2.0 and A^260^/_280_ between 1.8-2.2^50^. With the protocol described by Ziros et al., we were able to obtain RNA with A^260^/_280_ of 2.03±0.03 and A^260^/_230_ of 1.9±0.4 (Table 1). It should be noted that these ratios depend on the concentration of both RNA and contaminants – so trace amounts of a contaminant could significantly skew the purity ratio if the RNA concentration is also low. Typical contaminants that absorb at 230nm are phenol and guanidine thiocyanate found in lysis buffers. In high quantities these contaminants can inhibit PCR and give misleading expression results.

We also showed that cirrhotic slices remain viable in culture longer than slices from healthy animals. This may be due to adaptation in vivo to the hypoxic and poorly perfused cirrhotic liver^38-41^. This enhanced survival ex vivo adds to the benefits of PCLS from cirrhotic or fibrotic animals as a more appropriate model to test anti-fibrotic therapies than PCLS from healthy animals. Though some previous studies have used healthy PCLS to test antifibrotic therapies, this approach likely has limited application to clinically relevant liver fibrosis. The collagen deposition and increasing tissue stiffness in chronic liver fibrosis have a variety of effects on the liver. There is a crucial positive feedback loop between matrix stiffness and collagen deposition – increasing stiffness causes pro-fibrotic signaling, leading to collagen deposition and further increasing stiffness^22-25^ – which would not be established in PCLS from healthy animals. For this reason, treatments that can suppress fibrosis in soft tissues may be less effective in stiff tissues. Matrix stiffness also influences hepatocyte phenotype, with a stiffer matrix leading to loss of hepatocyte-specific functions ^19-21^. This would be important for functional outcome metrics and toxicology. These aspects are unlikely to sufficiently develop in healthy slices over just a few days in culture but can be captured with fibrosis induced in vivo.

PCLS are a promising platform for the future of antifibrotic therapy development. The interactions between different cell populations as well as interactions with their microenvironment is a crucial part of the response to disease, injury, and treatment. Especially in the setting of fibrosis, cell-matrix biomechanics plays an important regulatory role. Other methods that attempt to address this include organoids and 3D co-culture platforms that mimic parts of the in vivo liver^51-53^. PCLS is a relatively simple technique that directly captures all liver cell populations in their native architecture. PCLS also has the benefit of being able to be used with human liver biopsies – enabling new compounds to be tested on human tissue and potentially opening the possibility of personalized medicine applications.

Cirrhotic PCLS also offer several advantages over in vivo models. Cirrhosis takes several months to develop in mice and rats. With the PCLS model, each cirrhotic animal can provide dozens of PCLS which can be divided between many different experimental groups. This enables higher throughput testing of new drugs, while also reducing the number of required animals. Where appropriate, PCLS also provide the opportunity for highly controlled experiments as nearly identical serial slices can be tested under multiple conditions. Ultimately PCLS are not a replacement for in vivo testing – given the lack of circulating immune cells and inability to identify potential toxicity outside the liver, among other drawbacks – but they enable faster and easier screening of compounds in a realistic and meaningful ex vivo platform which can substantially reduce the number of animals required for drug development.

## Acknowledgements

RRA was partially supported by the Lancer Endowed Chair in Dermatology

## Author Contributions

Conceptualization: BDL, YW, JT, RRA. Formal analysis: BDL. Funding acquisition: RRA. Investigation: BDL, YW. Methodology: BDL, YW. Supervision: JT, RRA. Writing – original draft: BDL. Writing – review & editing: YW, JT, RRA.

## Data availability

The datasets generated during and/or analyzed during the current study are available from the corresponding author on reasonable request.

## Additional Information

The authors declare no competing interests.

