## Supplementary Material for "Analysis of culture and RNA isolation methods for precision-cut liver slices from cirrhotic rats"

**SUPPLEMENTAL INFORMATION**

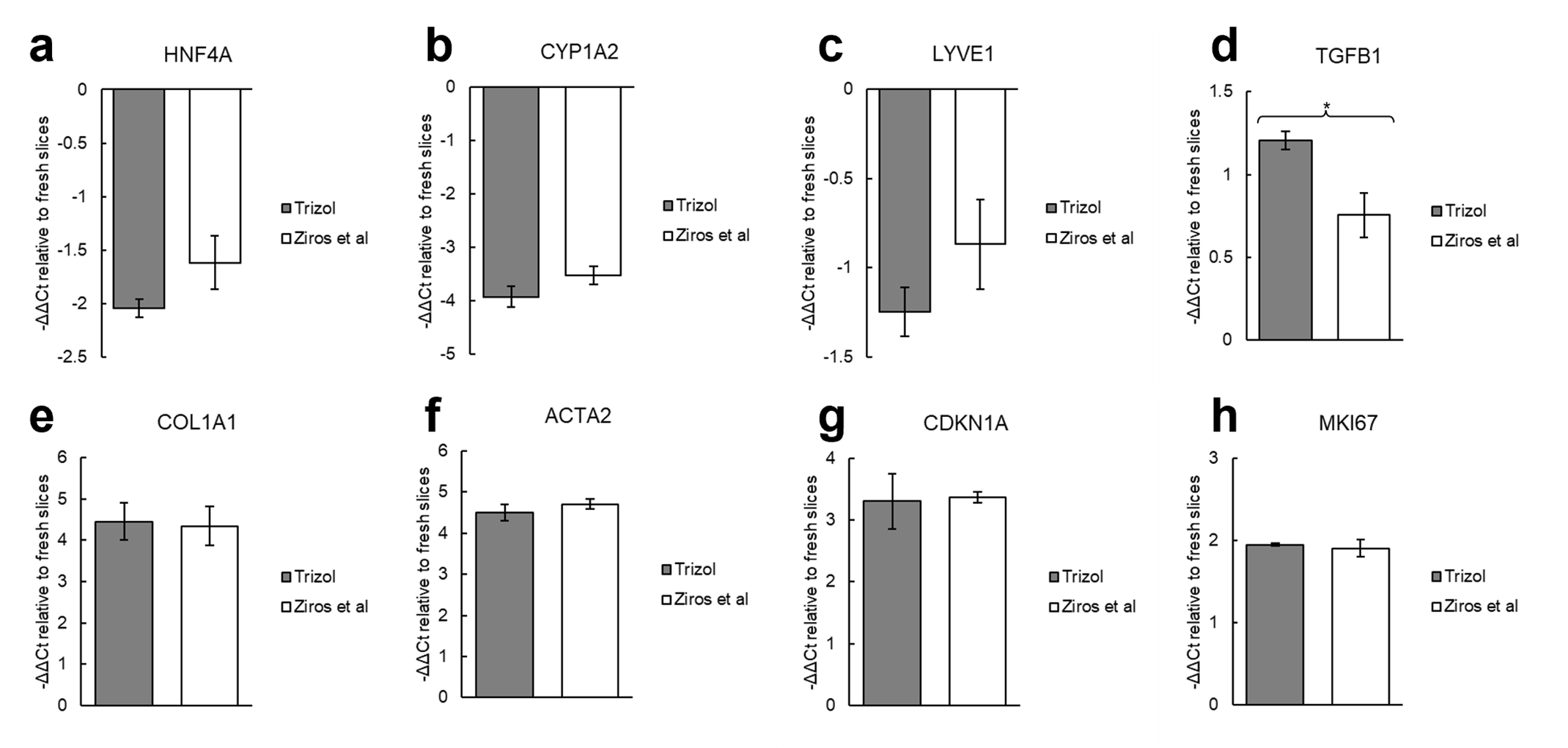

**Supplementary Figure 1: PCR results for RNA isolated with TRIzol protocol or Ziros et al. protocol from cirrhotic PCLS after 4 days in culture.** *p<0.05

| Time in Culture | Yield  (µg/mg tissue) | A$\frac{260}{280}$ | A$\frac{260}{230}$ |
| --- | --- | --- | --- |
| Fresh Slices | 6.3±1.8 | 2.08±0.02 | 2.1±0.1 |
| 4d | 2.3±0.9^*^ | 2.03±0.03^*^ | 1.9±0.4 |

**Supplementary Table 1: Yield and purity metrics for RNA isolated from fresh cirrhotic PCLS and after 4 days in culture.** RNA was isolated with the protocol described by Ziros et al. *p<0.05
